# Acute Effects of Prolonged Intermittent Theta Burst Stimulation on Cortical Excitability: A Randomized Crossover TMS-EEG Study

**DOI:** 10.64898/2026.11.27.769602

**Authors:** Sharie Xiao Wang, Wanda Man Wai Chau, Vera Wai Man Lam, Tim Tianze Lin, Roy Rongyue Zeng, Penny Ping Qin, Minxia Jin, Adam Weili Xia, Jack Jiaqi Zhang, Tifei Yuan, Yoshihiro Noda, Georg S. Kranz

**Affiliations:** Department of Rehabilitation Sciences, The Hong Kong Polytechnic University, Hong Kong, China; Shanghai YangZhi Rehabilitation Hospital (Shanghai Sunshine Rehabilitation Center), School of Medicine, Tongji University, Shanghai, China; Shanghai Key Laboratory of Mental Disorders Translational Research, Brain Health Institute, National Center for Mental Disorders, Shanghai Mental Health Center, Shanghai Jiao Tong University School of Medicine and School of Psychology, Shanghai, China; Japan Center for Advanced Neuromodulation, Seishinkai Kakoro Medical Clinic, Yokohama, Japan; Mental Health Research Center (MHRC), The Hong Kong Polytechnic University, Hong Kong, China; University Research Facility in Behavioral and Systems Neuroscience (UBSN), The Hong Kong Polytechnic University, Hong Kong, China

**Author notes:** **Correspondence to:** Dr. Georg S. Kranz, Department of Rehabilitation Sciences, The Hong Kong Polytechnic University Hong Kong, SAR, China.

**Keywords:** TMS-EEG, prolonged intermittent theta burst stimulation, Stanford Neuromodulation Therapy, theta oscillations, cortical reactivity, N100

## Abstract

**Background:** prolonged intermittent theta burst stimulation (piTBS), as used in Stanford Neuromodulation Therapy (SNT), achieves antidepressant efficacy, yet its acute single-session mechanisms remain elusive. EEG can capture online oscillatory dynamics during piTBS and pre-to post-stimulation changes in TMS-evoked cortical reactivity.

**Methods:** in a randomized, sham-controlled crossover trial, 40 healthy males completed two visits (≥ 7-day washout), receiving active or sham piTBS over the left dorsolateral prefrontal cortex. Inter-train EEG power was analyzed across three blocks of 20 trains in 37 participants. Pre- and post-stimulation measures of TMS-evoked potentials (TEPs), working memory and sustained attention were analyzed in all 40 participants.

**Results:** left frontal theta power followed different trajectories across blocks during active and sham piTBS (condition × block interaction, *p* = .014). During active piTBS, theta power was below baseline in blocks 1 and 2 and increased significantly from block 2 to block 3. Across conditions, alpha power was lower in blocks 2 and 3 than in block 1. Direct comparison of pre-to-post TEP changes between active and sham piTBS identified no significant clusters in the N45, P60, N100 and P180 component windows. Exploratory analyses within active piTBS identified a decrease in frontal N100 cluster amplitude and an increase in posterior N100 cluster amplitude. Neither behavioral outcome showed condition-dependent changes.

**Conclusion:** a single session of active piTBS modulated left frontal theta power between stimulation trains. Whether this early response is associated with cumulative plasticity or antidepressant outcomes following SNT warrants further investigation.

## 1. Introduction

Treatment-resistant depression (TRD) is a highly prevalent and disabling condition associated with substantial personal and economic burden [1,2]. In the STAR*D trial, an estimated 44% of patients did not achieve remission after two treatment steps [3]. Although conventional intermittent theta burst stimulation (iTBS) targeting the left dorsolateral prefrontal cortex (DLPFC) delivers 600 pulses per session and is FDA-cleared for adults with major depressive disorder following an inadequate response to antidepressant medication, treatment typically spans several weeks and clinical outcomes vary [4]. As an accelerated repetitive transcranial magnetic stimulation (rTMS) paradigm, the Stanford Neuromodulation Therapy (SNT) protocol employs a prolonged schedule of 1800 pulses per session (prolonged iTBS, piTBS) and ten sessions per day, demonstrating substantial efficacy, with response rates of approximately 70% at the one-month follow-up [5,6]. This accelerated response is hypothesized to be driven by a robust, cumulative induction of long-term potentiation (LTP)-like synaptic plasticity [7]. However, the immediate neurophysiological mechanisms driving this response remain elusive, and to what extent a single 1800-pulses piTBS session acutely modulates prefrontal reactivity is unknown, even as it serves as the fundamental unit of cumulative therapeutic efficacy.

Concurrent transcranial magnetic stimulation and electroencephalography (TMS-EEG) can characterize the acute neurophysiological effects of left DLPFC piTBS. Unlike motor cortex stimulation yielding motor evoked potentials (MEPs), DLPFC stimulation lacks a direct peripheral readout, limiting conventional TMS outcome measures [8]. In addition, neuroimaging approaches such as functional magnetic resonance imaging, positron emission tomography, and functional near-infrared spectroscopy do not provide the millisecond temporal resolution required to capture rapid stimulation-induced cortical dynamics [9]. By combining causal cortical perturbation with high-temporal-resolution recording, TMS-EEG directly assesses cortical responses relative to stimulation [10]. Single-pulse TMS-EEG can be used to derive TMS-evoked potentials (TEPs), which provide time-locked measures of cortical reactivity to detect pre-to post-stimulation changes [8]. TEPs comprise reproducible components, including N45, P60, N100, and P180, which have been studied in relation to cortical excitation and inhibition [11–13]. The N100 is particularly robust and highly reproducible across individuals and sessions [12,14,15]. Consequently, TMS-EEG has been widely used to probe stimulation-induced changes in prefrontal circuits [16–19].

Prior TMS-EEG studies have typically focused on changes in TEPs from before to after stimulation, giving comparatively little attention to neural activity unfolding during iTBS. This distinction matters because iTBS is a patterned stimulation protocol thought to engage rhythm-dependent plasticity mechanisms [20]. EEG recorded between iTBS trains can characterize frequency-specific changes in ongoing oscillatory activity. TEPs measured before and after the session could provide a complementary measure of changes in cortical reactivity. Accordingly, the present sham-controlled crossover study employed TMS-EEG to investigate the acute neurophysiological effects of a single piTBS session delivered to the left DLPFC in healthy participants. Studying healthy individuals reduced the influence of illness burden, medication, and clinical heterogeneity. We quantified EEG spectral power between piTBS trains to characterize online oscillatory dynamics, assessed changes in TEPs from before to after stimulation to evaluate modulation of cortical reactivity, and examined cognitive performance to determine accompanying behavioral effects. We hypothesized that, relative to sham, active piTBS would modulate frequency-specific online oscillations, alter TEP indices of cortical reactivity, particularly the N100 component, and produce corresponding improvements in cognitive performance.

## 2. Methodology

### 2.1 Study design and participants

This study was a two-visit, randomized, participant-blind, controlled crossover trial conducted at The Hong Kong Polytechnic University. Forty-six right-handed healthy male adults aged 20 to 30 years were recruited via campus posters from April 2025 to November 2025. These eligibility restrictions were applied to reduce sample heterogeneity and minimize potential sex- and age-related variability in cortical excitability [21–23]. Exclusion criteria were as follows: (1) a current or past diagnosis of a psychiatric or neurological disorder, or severe internal medical illness; (2) any contraindication to TMS or EEG; (3) a family history of psychiatric disorders in first-degree relatives; (4) substance abuse within the past 3 months; (5) current or recent use of any illicit drugs; and (6) neuroscience or medicine background.

This study was conducted in accordance with the Declaration of Helsinki and was approved by the Institutional Review Board of the Hong Kong Polytechnic University (HSEARS20241211006). This study was pre-registered on Open Science Framework and can be accessed via the following anonymized link for peer review: https://osf.io/fphr2/overview?view_only=cdfc84e265324f58b141e0bae7c0e658. All participants provided written informed consent prior to participation.

### 2.2 Study procedure and randomization

Participants completed two visits in a randomized and counterbalanced order separated by at least 7 days [24,25]. Randomization was generated using a two-step stratified block procedure (RANDOM.ORG) [26], allocating participants 1:1 to active-sham or sham-active piTBS sequences, with TMS-EEG trial order further stratified within each allocation block. Six participants withdrew before completing both visits; subsequently enrolled participants were assigned to the corresponding unfilled slots until 40 completers were reached.

Prior to visit 1, eligible participants were screened using a standardized form. On arrival, written informed consent was obtained and RMT was determined as described in *Section 2.3.2*. Pre-stimulation assessments then included a 3-back task, Sustained Attention to Response Task (SART), and a TMS-EEG session comprising 100 active and 100 sham single-pulse TMS (spTMS) trials. The assigned TMS-EEG trial order remained constant across all pre- and post-stimulation assessments for both visits. Participants then received active or sham left DLPFC piTBS with concurrent EEG recording, followed by identical post-stimulation assessments. Visit 2 followed the same procedure with the alternate condition of piTBS (Figure 1A).

**Figure 1.**
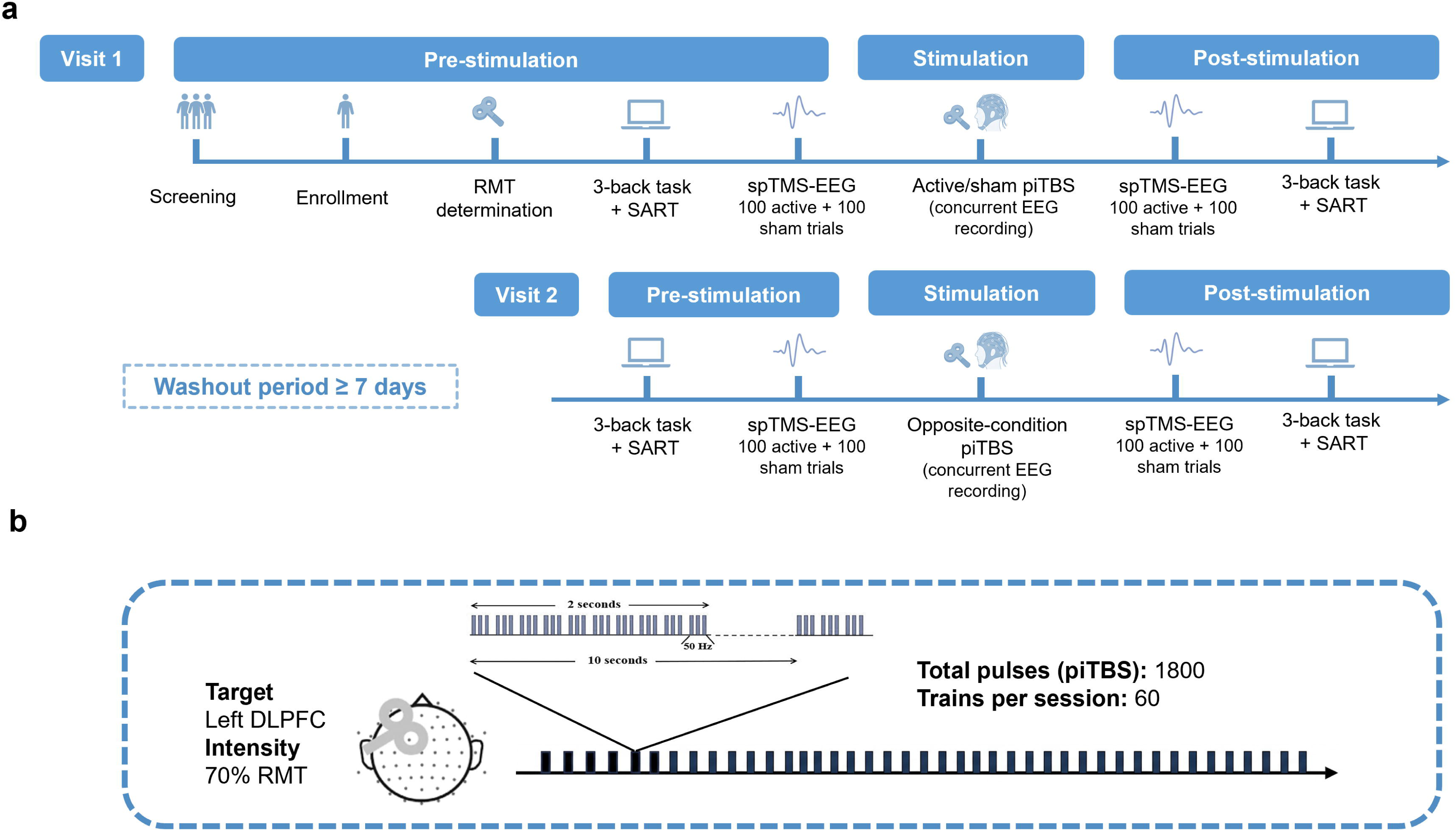
Experimental design and piTBS protocol. (A) Schematic overview of the experimental timeline and crossover design. The flowchart illustrates the sequence of procedures across the two visits, separated by a minimum 7-day washout period. (B) Illustration of the piTBS target and stimulation parameters. PiTBS was delivered over the left dorsolateral prefrontal cortex (DLPFC) at 70% of the resting motor threshold (RMT). Each session consisted of 60 trains for a total of 1800 pulses.

### 2.3 Transcranial magnetic stimulation

#### 2.3.1 Prolonged intermittent theta burst stimulation

The piTBS protocol [5] comprised 1800 pulses (10-burst trains at 50 Hz, 2 s train duration, 60 trains, 8 s inter-train interval; Figure 1B). Stimulation was delivered to left DLPFC at 70% RMT using a MagVenture Cool-B65 A/P figure-of-eight coil connected to a MagPro X100 stimulator (MagVenture, Denmark), with the active face used for active stimulation and the reverse face for sham. Functional evidence indicates that prefrontal iTBS at 70% RMT is sufficient to induce robust cortical activation and modulate executive function [27], while maximizing participant tolerability by minimizing scalp pain and facial muscle twitching [28]. The target was localized via neuronavigation system (LOCALITE, Germany) based on MNI coordinates (x = −38, y = 44, z = 26; Brodmann area, BA46) [29], with the coil positioned above the EEG cap at a 45° angle relative to the mid-sagittal line.

#### 2.3.2 Single-pulse TMS

Biphasic spTMS for RMT determination used a MagVenture MCF-B65 figure-of-eight coil. RMT was defined as the minimum intensity eliciting MEPs with a peak-to-peak amplitude of at least 50 µV in the relaxed right first dorsal interosseous muscle in 5 of 10 consecutive trials, achieved by stimulating the optimal hotspot of the left primary motor cortex (M1). The hotspot over the left M1 corresponding to the contralateral hand muscles was manually identified following the procedures described by Ortega-Robles et al. and Rossini et al. [30,31].

For the TMS-EEG, 100 single pulses were delivered per spTMS condition at 100% RMT with a 5 s inter-stimulus interval (±10% jitter). The pulse number [19] and stimulation intensity [18] were selected based on previous protocols investigating this region. The target and the coil configuration matched *Section 2.3.1*.

### 2.4 Electroencephalography

EEG data were acquired using a 64-channel TMS-compatible EEG cap (Easycap, Germany) and DC-coupled amplifier (SynAmps2, USA) at 5 kHz. The FCz and AFz served as recording reference and ground. Impedances were kept below 5 kΩ. Throughout EEG recording, participants remained in a seated position with their eyes closed and were instructed to stay awake. During spTMS-EEG blocks, white noise was delivered via intra-auricular earphones to minimize auditory artifacts. The sound level was individually adjusted until the spTMS clicks were masked by the white noise.

EEG data recorded during piTBS were preprocessed using custom scripts in MATLAB R2024b (MathWorks, USA) and EEGLAB [32] (resampled to 1,000 Hz, 1 Hz high-pass, 50 Hz notch, epochs 1,000–7,000 ms post-train, band-pass 1–100 Hz, common average reference) [33] (Figure 2B). SpTMS-EEG data were preprocessed using the AARATEP pipeline (v2) [34] in MATLAB R2024b (epochs −800 to +1,000 ms, pulse artifact removal, high-pass >1 Hz, bad channel reconstruction, ICLabel ICA, spatial noise suppression, low-pass <100 Hz; Figure 2A). Pipeline source code is available at <u>github.com/chriscline/AARATEPPipeline</u>. Full details are in *Supplementary Methods S1*.

**Figure 2.**
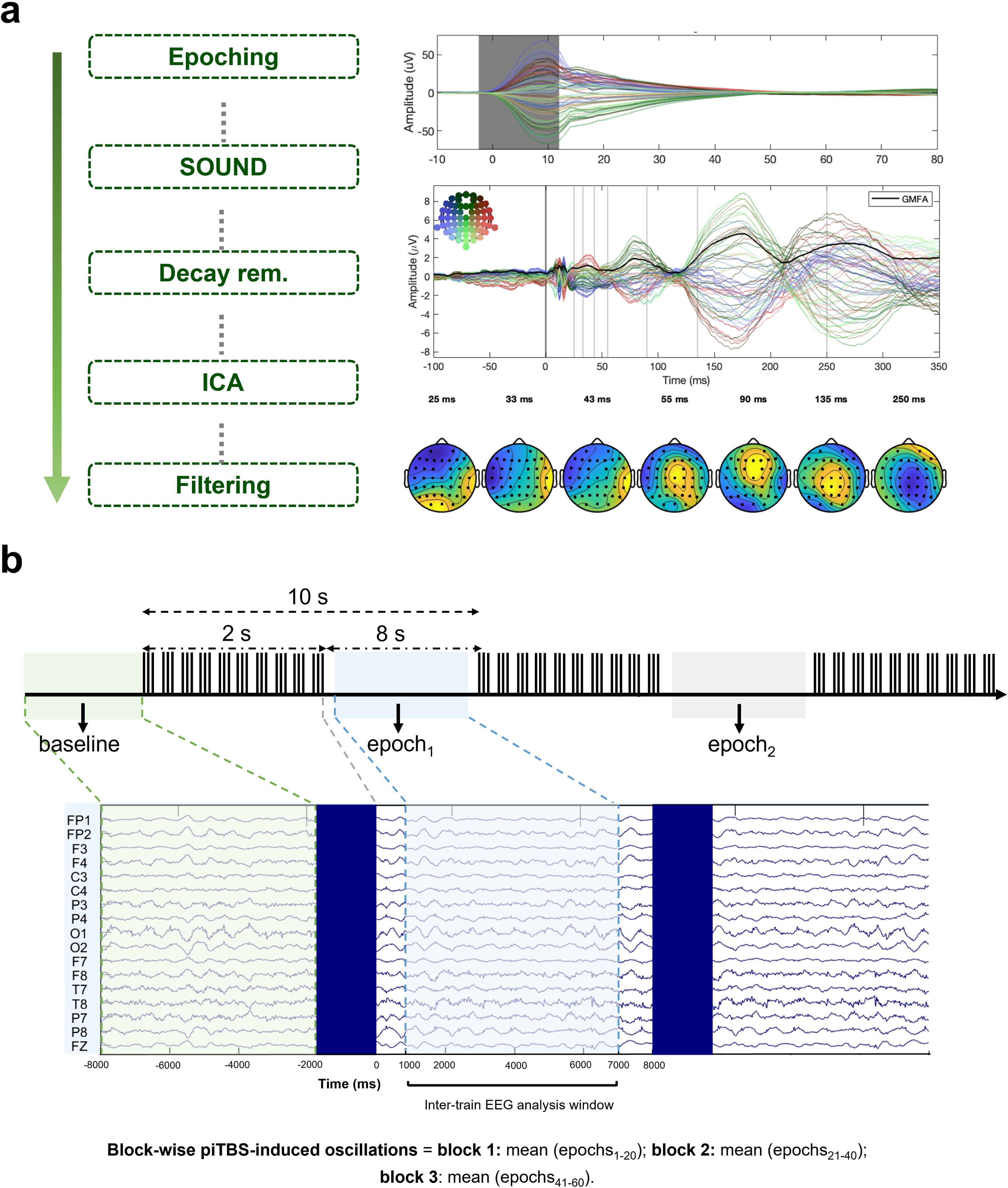
spTMS-EEG data preprocessing and frequency-domain analysis pipeline for piTBS-induced oscillations. (A) Selected steps of the automated preprocessing pipeline described by Cline et al. [34] are shown on the left. Artifact amplitude was quantified before artifact removal (SOUND) (top right). Example butterfly plots (100 trials, N = 1 subject) (bottom right). (B) Schematic illustration of the segmentation procedure for frequency-domain analysis of inter-train oscillatory activity. Time 0 represents the end of each piTBS train. Spectral power was estimated from epochs 1,000–7,000 ms after each train and expressed in dB relative to the 6-s baseline preceding the first pulse of the session. Block-wise oscillatory activity was calculated as the mean of epochs 1–20, 21–40, and 41–60 for blocks 1, 2, and 3, respectively. Abbreviations: piTBS, prolonged intermittent theta burst stimulation; EEG, electroencephalography; spTMS, single-pulse transcranial magnetic stimulation.

### 2.5 Cognitive function assessments

The cognitive evaluation paradigm consisted of a visual 3-back task [35] and the SART [36], with a randomized task presentation order. The visual 3-back task was used to evaluate working memory, given its established sensitivity to DLPFC-dependent cognitive processes [37]. Each block contained 30 trials (10 targets, 20 non-targets). Performance was indexed by d′ = Z (hit rate) – Z (false alarm rate). The SART was administered to provide a complementary assessment of sustained attention and response inhibition as indices of prefrontal-dependent executive function [38]. Participants responded to digit stimuli (1–9) and withheld responses to the target digit (3); Go: No-Go ratio was 8:1. Performance was indexed by commission error rate, defined as the proportion of No-Go trials on which participants responded. Full details are in *Supplementary Methods S2 and S3*.

### 2.6 Statistical analysis

PSD-derived alpha- and theta-band power values in a left frontal region of interest, defined a priori as the mean of electrodes F3 and AF3, were analyzed using separate repeated-measures ANOVAs (rmANOVAs) with condition (active vs. sham piTBS) and block (block 1, block 2, block 3) as within-subject factors. Significant main effects or interactions were followed by Bonferroni-corrected post hoc paired-samples t-tests. One-sample t-tests against 0 dB were also conducted to assess deviations from baseline, with Bonferroni correction applied for multiple comparisons. TEPs were analyzed using paired spatiotemporal cluster-based permutation tests across all scalp electrodes within four predefined component windows (N45 [30–50 ms], P60 [50–70 ms], N100 [95–125 ms] and P180 [160–200 ms]). We directly compared post-minus-pre TEP changes between active and sham piTBS. Exploratory analyses compared pre- and post-stimulation TEPs separately within each condition. Each test used 5,000 permutations, and two-sided cluster *p* values were Bonferroni-adjusted across the four windows. Mean amplitudes within clusters identified in the exploratory analyses were analyzed using separate 2 × 2 rmANOVAs with condition (active vs. sham piTBS) and time (pre-stimulation vs. post-stimulation) as within-subject factors. Separate linear mixed-effects models were fitted for 3-back discriminability (d′) and SART commission error rate. Each model included time (pre-vs. post-stimulation), condition (active vs. sham piTBS), the condition × time interaction and visit order (visit 1 vs. visit 2) as fixed effects. Age and years of education were included as covariates, and participant was included as a random intercept. All *t*-tests were two-sided. Statistical significance was assessed at *p* < .05. Full details are in *Supplementary Methods S4*.

## 3. Results

### 3.1 Participants

Of the 46 participants who provided informed consent, 40 (mean age = 23.10 ± 2.12 years; range = 20–28 years; mean years of education = 19.56 ± 2.05 years; range = 14–23 years) completed the study. Six participants withdrew, including one for personal reasons and five whose RMT exceeded the maximum stimulator output, preventing threshold determination. Behavioral and spTMS-EEG data from all 40 participants were analyzed. For the piTBS-EEG analyses, 37 datasets were included (mean age = 22.76 ± 2.09 years, range = 20–28 years; mean years of education = 19.28 ± 2.17 years, range = 14–23 years) after excluding three datasets due to poor data quality.

### 3.2 piTBS-induced oscillatory responses

Left frontal theta-band power showed significant main effects of condition and block and a significant condition × block interaction (Figure 3A and Table 1). After Bonferroni correction, theta power during active piTBS was higher in block 3 than in block 2. Neither block 2 nor block 3 differed significantly from block 1. No pairwise differences reached significance in the sham piTBS condition (Figure 3B and Table 1). Relative to the 0 dB baseline, theta power during active piTBS was significantly lower in blocks 1 and 2 (adjusted *p* = .019 and *p* < .001, respectively). Block 3 in the active condition and all sham blocks did not differ significantly from baseline (Figure 3A).

**Figure 3.**
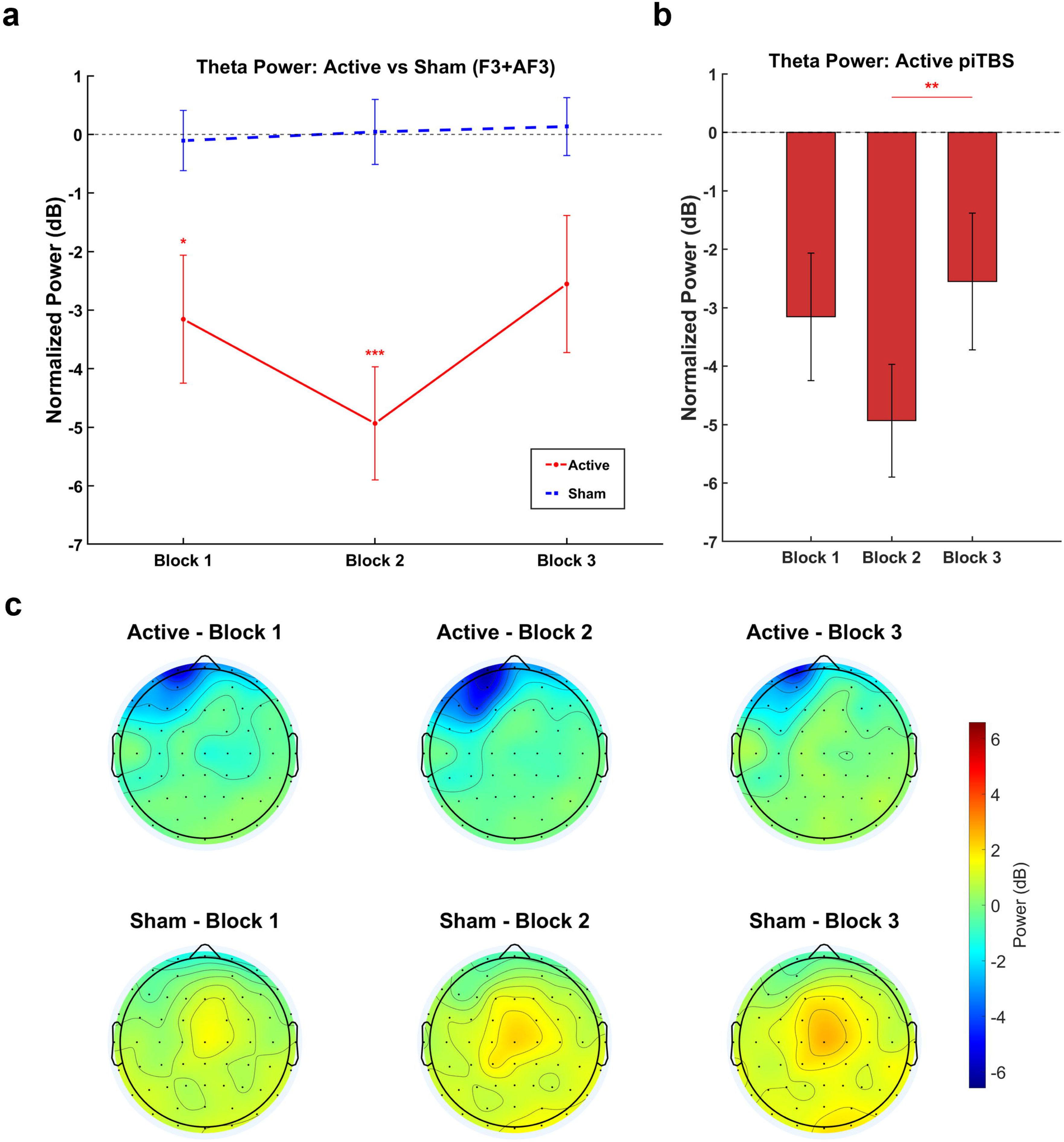
Theta-band power in the left frontal ROI (mean of F3 and AF3) across blocks under active and sham piTBS. (A) Theta-band normalized power (dB) across blocks 1–3. The red solid line denotes the active condition, and the blue dashed line denotes the sham condition. The gray dashed horizontal line indicates baseline (0 dB). Asterisks indicate significant Bonferroni-adjusted comparisons with the 0 dB baseline. Error bars represent SEM. (B) Theta-band normalized power (dB) in the active condition across blocks 1–3. Bars represent mean values, and error bars represent SEM. The horizontal line and asterisks indicate the significant Bonferroni-adjusted comparison between blocks 2 and 3. (C) Scalp topographies of theta-band power across Blocks 1–3 in the active and sham conditions. * *p* < .05, ** *p* < .01, *** *p* < .001.

**Table 1.** Statistical results for spectral power, TMS-evoked potentials and behavioral outcomes.

| **A. Spectral power** |  |  |  |
| **Measure** | **Effect or comparison** | **Test statistics** | ***p*-value** |
|---|---|---|---|
| **Left Frontal Theta-band Power** | Condition main effect | *F* <sub>(1,36)</sub> = 13.41 | < .001 |
|  | Block main effect | *F* <sub>(2,72)</sub> = 6.08 | .004 |
|  | Condition × Block interaction | *F* <sub>(2,72)</sub> = 4.50 | .014 |
|  | *Active piTBS:* block 1 – block 2 | *t* <sub>(36)</sub> = 2.27 | .088 |
|  | *Active piTBS:* block 2 – block 3 | *t* <sub>(36)</sub> = -3.25 | .007 |
|  | *Active piTBS:* block 1 – block 3 | *t* <sub>(36)</sub> = -1.49 | .436 |
|  | *Sham piTBS:* block 1 – block 2 | *t* <sub>(36)</sub> = -0.33 | 1.000 |
|  | *Sham piTBS:* block 2 – block 3 | *t* <sub>(36)</sub> = -0.28 | 1.000 |
|  | *Sham piTBS:* block 1 – block 3 | *t* <sub>(36)</sub> = -0.60 | 1.000 |
| **Left Frontal Alpha-band Power** | Condition main effect | *F* <sub>(1,36)</sub> = 0.85 | .364 |
|  | Block main effect | *F* <sub>(2,72)</sub> = 22.02 | < .001 |
|  | Condition × Block interaction | *F* <sub>(2,72)</sub> = 2.23 | .115 |
|  | Averaged across piTBS conditions: block 1 – block 2 | *t* <sub>(36)</sub> = 6.20 | < .001 |
|  | Averaged across piTBS conditions: block 2 – block 3 | *t* <sub>(36)</sub> = 0.73 | 1.000 |
|  | Averaged across piTBS conditions: block 1 – block 3 | *t* <sub>(36)</sub> = 5.78 | < .001 |
| **B. TMS-evoked potentials** |  |  |  |
| **Analysis** | **Effect or comparison** | **Test statistic/cluster extent** | ***p*-value** |
| **Direct ΔTEP comparison** | ΔTEP after active piTBS vs. ΔTEP after sham piTBS | No significant clusters |  |
| **Exploratory within-condition cluster tests** | *Active piTBS:* Post – Pre, negative frontal N100 cluster | 95–122 ms | .027 |
|  | *Active piTBS:* Post – Pre, positive posterior N100 cluster | 95–114 ms | .030 |
| **Frontal N100 cluster amplitude** | Time main effect | *F* <sub>(1,39)</sub> = 10.77 | .002 |
|  | Condition main effect | *F* <sub>(1,39)</sub> = 0.08 | .784 |
|  | Condition × Time interaction | *F* <sub>(1,39)</sub> = 0.63 | .431 |
|  | *Active piTBS:* Post – Pre (Simple effect) | *t* <sub>(39)</sub> = -3.04 | .017 |
| **Posterior N100 cluster amplitude** | Time main effect | *F* <sub>(1,39)</sub> = 10.86 | .002 |
|  | Condition main effect | *F* <sub>(1,39)</sub> = 0.07 | .790 |
|  | Condition × Time interaction | *F* <sub>(1,39)</sub> = 0.44 | .509 |
|  | *Active piTBS:* Post – Pre (Simple effect) | *t* <sub>(39)</sub> = 3.33 | .008 |
| **C. Behavioral outcomes** |  |  |  |
| **Measure** | **Fixed effect** | **Test statistic** | ***p*-value** |
| **3-back discriminability (*d′*)** | Time | *F* <sub>(1,116)</sub> = 7.03 | .009 |
|  | Condition | *F* <sub>(1,116)</sub> = 0.08 | .771 |
|  | Condition × Time | *F* <sub>(1,116)</sub> = 0.53 | .468 |
|  | Visit order | *F* <sub>(1,116)</sub> = 9.02 | .003 |
| **SART commission error rate** | Time | *F* <sub>(1,116)</sub> = 0.01 | .904 |
|  | Condition | *F* <sub>(1,116)</sub> = 0.31 | .579 |
|  | Condition × Time | *F* <sub>(1,116)</sub> = 0.05 | .828 |
|  | Visit order | *F* <sub>(1,116)</sub> = 1.18 | .279 |

*Note.* PSD = power spectral density. TEPs = TMS-evoked potentials. piTBS = prolonged intermittent theta burst stimulation. SART = Sustained Attention to Response Task. Condition refers to active versus sham piTBS. Blocks 1, 2 and 3 correspond to stimulation trains 1–20, 21–40 and 41–60, respectively. Time refers to the pre- and post-piTBS assessments. ΔTEP denotes the post-minus-pre change in the TEP waveform. Visit order refers to the first versus second study visit. Bonferroni adjustment was applied to *p* values for pairwise block comparisons, TEP cluster tests and simple-effect comparisons.

Left frontal alpha-band power showed a significant main effect of block. Neither the condition main effect nor the condition × block interaction was significant (Figure 4A and Table 1). Averaged across active and sham piTBS, alpha-band power was lower in blocks 2 and 3 than in block 1 after Bonferroni correction. Blocks 2 and 3 did not differ significantly (Table 1). Across conditions, alpha-band power was significantly below the 0 dB baseline in blocks 2 and 3 (both adjusted *p* < .001). Block 1 did not differ significantly from baseline (adjusted *p* = .485).

**Figure 4.**
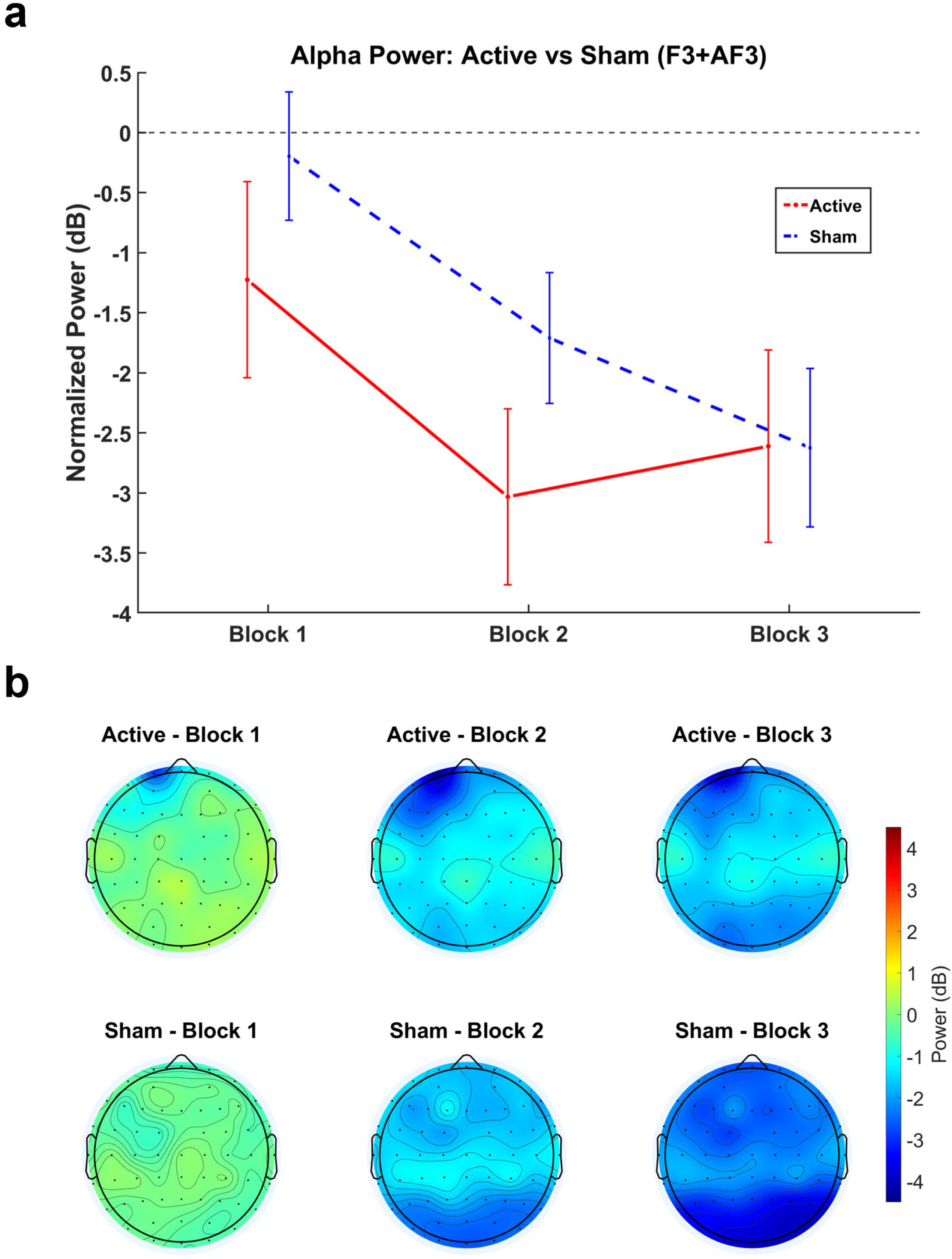
Alpha-band power in the left frontal ROI (mean of F3 and AF3) across blocks under active and sham piTBS. (A) Alpha-band normalized power (dB) across blocks 1–3. The red solid line denotes the active condition, and the blue dashed line denotes the sham condition. The gray dashed horizontal line indicates baseline (0 dB). Error bars represent SEM. (B) Scalp topographies of baseline-normalized alpha-band power (dB) across blocks 1–3 in the active and sham conditions.

### 3.3 TMS-evoked potentials

After preprocessing, TEP waveforms recorded with active single-pulse TMS showed the expected morphology of DLPFC-evoked responses, whereas sham single-pulse TMS waveforms showed no comparable pattern (Figure 5A). We first compared post-minus-pre TEP changes between active and sham piTBS using paired spatiotemporal cluster-based permutation tests across all scalp electrodes. No significant clusters were detected in any of the four predefined windows (N45, 30–50 ms, P60, 50–70 ms, N100, 95–125 ms, and P180, 160–200 ms).

**Figure 5.**
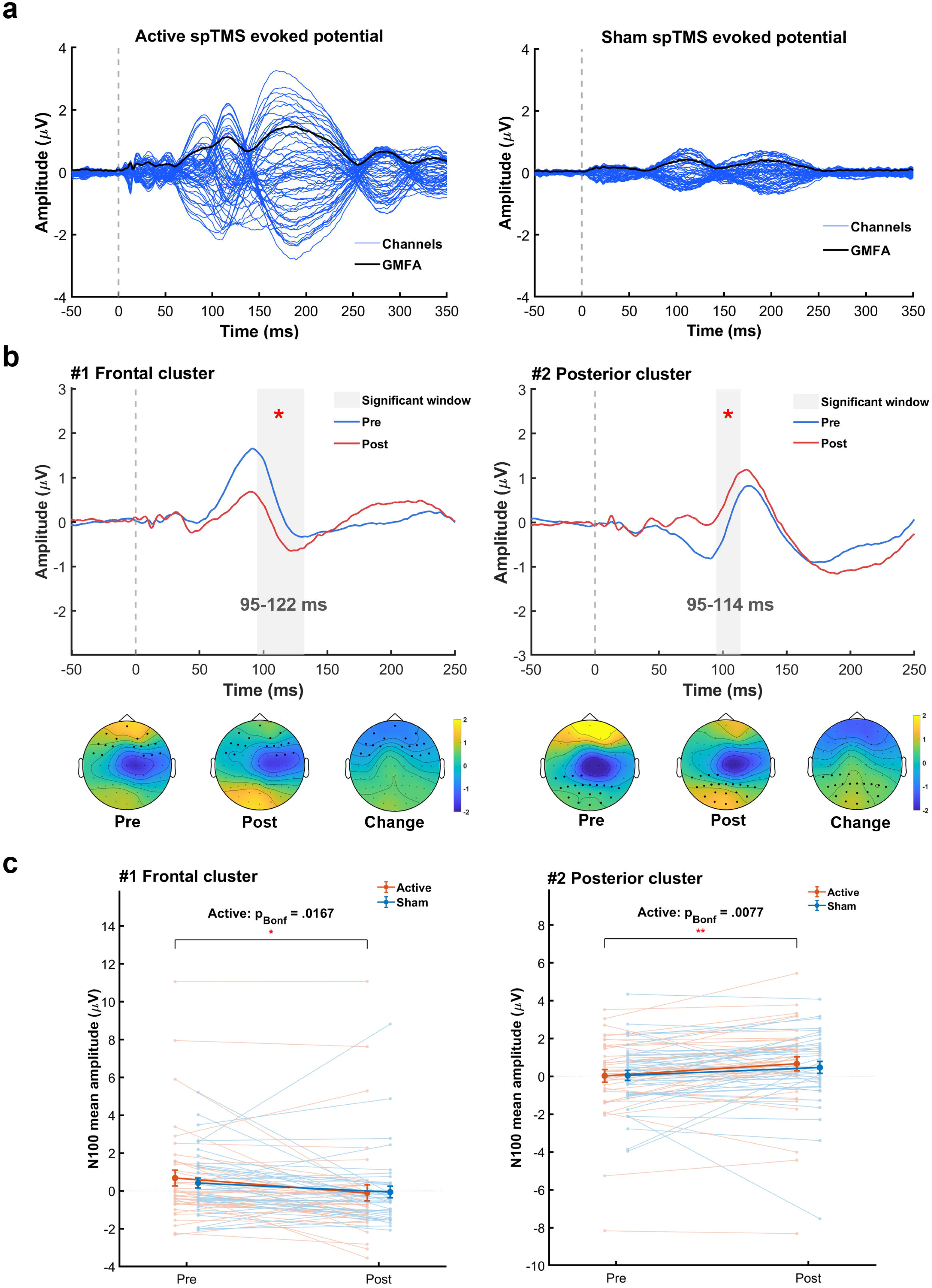
TEP waveforms and cluster-level changes following active and sham piTBS. (A) Butterfly plots of preprocessed TMS-evoked potentials (TEPs) elicited by active (left) and sham (right) single-pulse TMS. Blue lines indicate EEG channels, and the black line indicates global mean field amplitude (GMFA). The dashed vertical line marks TMS onset (0 ms). (B) Exploratory cluster-based permutation comparisons of post-versus pre-stimulation TEPs within the active piTBS condition identified a negative frontal cluster (#1, 95–122 ms, adjusted *p* = .027) and a positive posterior cluster (#2, 95–114 ms, adjusted *p* = .030). Blue and red lines represent pre- and post-piTBS waveforms, respectively. Shading marks the temporal extent of each cluster. Asterisks indicate cluster-level significance after Bonferroni adjustment across the four component windows. Topographic maps show pre-stimulation, post-stimulation, and post-minus-pre amplitudes averaged over the corresponding cluster interval. (C) Mean amplitudes of the frontal (left) and posterior (right) clusters before and after active and sham piTBS. Thin lines connect values from the same participant, and thick lines represent group means. Error bars show SEM. Brackets indicate Bonferroni-adjusted pre–post comparisons within the active piTBS condition. * *p* < .05, ** *p* < .01.

Exploratory within-condition analyses identified two N100 clusters after active piTBS. A negative frontal cluster extended from 95 to 122 ms, and a positive posterior cluster extended from 95 to 114 ms. Both remained significant after Bonferroni correction across the four component windows. No significant N100 clusters were detected after sham piTBS, and neither condition showed significant clusters in the other three windows (Figure 5B and Table 1).

Exploratory rmANOVAs on mean amplitudes within the two N100 clusters showed a significant main effect of time in both clusters. Neither cluster showed a significant main effect of condition or a condition × time interaction. Within-condition comparisons showed a pre-to-post decrease in frontal cluster amplitude and an increase in posterior cluster amplitude after active piTBS. Neither pre-to-post change reached significance after sham piTBS (Figure 5C and Table 1).

### 3.4 Cognitive tasks

The analysis of 3-back discriminability (d′), a signal detection measure of working memory performance, showed significant main effects of time and visit order. Discriminability was higher after stimulation than before stimulation and higher at visit 2 than at visit 1. Neither the main effect of condition nor the condition × time interaction was significant (Figure S1A and Table 1). For sustained attention and response inhibition on the SART, commission error rate, defined as the proportion of No-Go trials on which participants responded, showed no significant main effects of time, condition, or visit order. The condition × time interaction was also not significant (Figure S1B and Table 1). Taken together, these findings indicate that 3-back discriminability improved over time and was higher at visit 2 than at visit 1, consistent with a practice effect, whereas commission error rate on the SART remained stable across time, condition, and visit order.

## 4. Discussion

The present study used TMS-EEG to investigate the acute effects of a single piTBS session delivered to the left DLPFC in healthy participants, capturing both oscillatory dynamics between stimulation trains and pre-to-post changes in TEPs. Three principal findings emerged. First, left frontal theta-band power was lower overall during active than during sham piTBS, and its trajectory across blocks differed between conditions. During active piTBS, mean theta power followed a non-monotonic trajectory, declining to a nadir in block 2 before increasing significantly from block 2 to block 3. No pairwise block differences reached significance during sham piTBS. Alpha-band power declined mainly from block 1 to block 2 and was below baseline in blocks 2 and 3, with no clear evidence that this pattern differed by condition. Second, the direct comparison of pre-to-post TEP changes between active and sham piTBS identified no significant clusters in the four predefined component windows. Exploratory comparisons within the active piTBS condition showed a pre-to-post decrease in frontal N100 cluster amplitude and an increase in posterior N100 cluster amplitude. Third, we found no evidence that a single session of active piTBS improved working memory or sustained attention relative to sham.

The non-monotonic trajectory of frontal theta power suggests that the oscillatory response to active piTBS evolved within a single session. Theta-burst protocols were inspired by animal studies showing that stimulation patterned at theta frequency can induce hippocampal long-term potentiation [39–41]. Subsequent work has linked theta coupling in hippocampal–prefrontal circuits to synaptic plasticity in animals [42] and shown that the theta phase at which prefrontal rTMS is delivered can influence evoked responses in humans [43]. In our study, ongoing frontal theta power between stimulation trains was below the pre-stimulation baseline in blocks 1 and 2. A similar reduction in left prefrontal theta power has been reported in resting EEG after left DLPFC iTBS [44]. A separate study of left DLPFC iTBS found a reduction in theta power elicited by single-pulse TMS after stimulation, in an analysis across 600-, 1200-, and 1800-pulse protocols [45]. The significant rise from block 2 to block 3 indicates that the early reduction became less pronounced as stimulation continued. Human intracranial recordings have also shown that responses evoked by direct electrical theta-burst stimulation can change across successive trains [46]. The late rise in inter-train theta power may reflect adaptation to repeated stimulation, potentially involving homeostatic adjustment [47,48]. Studies of the human motor cortex have also shown that the after-effects of iTBS can vary with pulse count and the spacing of repeated stimulation blocks [49,50]. In one study, a 1,200-pulse iTBS protocol reduced motor-evoked potential amplitudes relative to sham, and a 1,800-pulse protocol showed no significant difference from sham [49]. This dependence of motor cortical after-effects on stimulation parameters provides a rationale for considering whether prior stimulation within the session also shaped the inter-train theta response.

Frontal alpha power declined mainly from block 1 to block 2 and was below the pre-stimulation baseline in blocks 2 and 3. We found no clear evidence that this trajectory differed between active and sham piTBS. In eyes-closed resting EEG, alpha power has been reported to decrease both after caffeine intake, alongside an increase in skin conductance, and after partial sleep restriction [51,52]. Resting-state magnetoencephalography recordings have similarly shown lower relative upper-alpha power during drowsy eyes-closed periods than during alert eyes-closed periods [53]. Fluctuations in alertness during the eyes-closed recording may therefore have contributed to the observed decline in inter-train alpha power.

The direct comparison of pre-to-post changes between active and sham piTBS identified no significant clusters in the N100 window. Exploratory within-condition analyses showed increased frontal negativity and reduced posterior negativity in the N100 time window from pre-to post-stimulation in the active piTBS. Both clusters showed a main effect of time, with no significant condition × time interaction. The N100 can be measured reproducibly after prefrontal TMS [54]. Du et al. found that the frontal glutamate-to-GABA ratio measured by magnetic resonance spectroscopy accounted for approximately 46% of the between-participant variance in frontal N100 amplitude in their sample [55]. Premoli et al. reported that baclofen, a GABA_B_ receptor agonist, increased N100 amplitude following motor cortex TMS [10]. These findings provide a rationale for investigating a possible GABAergic contribution to the frontal shift observed here. Previous prefrontal studies have reported increased N100 amplitude after one or two blocks of standard iTBS [56], decreased amplitude after individualized iTBS [57], and no significant N100 effect in a sham-controlled comparison of prefrontal TBS protocols [58]. The frontal and posterior changes warrant independent confirmation because both clusters were identified from the pre-to-post contrast within the active piTBS.

Working memory performance (3-back d′) improved from pre-to post-intervention across both conditions and was higher at the second visit than at the first. The visit effect is consistent with practice from repeated task exposure [59]. The pre-to-post improvement did not differ significantly between active and sham piTBS. Similarly, SART commission errors showed no significant effects of time, stimulation condition, or visit, and no significant time × condition interaction. Using a 1800 pulse piTBS protocol targeting the left DLPFC, Xu et al. likewise found no behavioral benefit relative to sham on the 2-back or Stroop task [60]. Feng et al. reported pre-to-post improvements in 2-back and 3-back accuracy without significant time × condition interactions, a pattern they interpreted as consistent with practice effects [61]. In our study, a single session of active piTBS provided no clear additional benefit over sham on the working memory or response inhibition measures.

Taken together, our findings show that frontal theta power between stimulation trains followed a condition-dependent trajectory within a single piTBS session. The direct comparison of pre-to-post TEP changes between active and sham piTBS identified no significant clusters, and neither behavioral outcome showed a significant condition × time interaction. Inter-train theta power characterized ongoing oscillatory activity during piTBS, while TEPs assessed cortical responses to spTMS before and after the session. Small or variable TEP after-effects may have been difficult to detect, given the limited repeatability of changes in prefrontal N100 amplitude following a single iTBS session [54]. Another possibility is that prefrontal reactivity changed transiently during piTBS and that the change had diminished by the post-stimulation TEP assessment. In the human motor cortex, a priming block of iTBS attenuated the effect of a subsequent block, consistent with homeostatic metaplasticity [62]. This finding raises the possibility that homeostatic regulation also shapes the time course of prefrontal reactivity during piTBS. The inter-train theta modulation observed here may represent an acute response that could build across repeated sessions and be associated with later changes in cortical reactivity or behavior. In TRD patients, longitudinal TMS-EEG recordings showed a reduction in early TMS-evoked responses at the left DLPFC target by the third day of active SNT [63]. An exploratory analysis of a separate two-week prefrontal piTBS trial linked changes in frontal resting-state theta–alpha amplitude modulation to symptom improvement [64]. Associations with clinical outcomes may depend on the EEG measure and stimulation protocol. Baseline task-modulated frontal theta predicted antidepressant response to 10-Hz rTMS, with no significant predictive association in the piTBS group [65]. Tracking inter-train theta from the first session across a full course of piTBS could establish how this acute response evolves with repeated stimulation and whether the first-session trajectory is associated with subsequent changes in cortical reactivity or depressive symptoms.

TMS-EEG allowed us to track how frontal theta power evolved between stimulation trains within a single piTBS session. Several limitations warrant consideration. First, the sample was restricted to healthy young men to reduce potential age- and sex-related variability, which limits generalizability to broader populations, including patients with TRD [21–23]. Second, examining only a single piTBS session leaves open how acute changes evolve across a multiday course. Third, the proposed contributions of GABAergic and glutamatergic signaling remain indirect and could be investigated using pharmacological approaches or magnetic resonance spectroscopy. Future studies in patients with TRD could test whether the inter-train theta trajectory observed here is reproducible and whether it predicts symptom improvement over a full treatment course. If this association is confirmed, subsequent studies could evaluate whether the measure can inform the selection of pulse dose or session frequency.

## 5. Conclusion

This study characterized acute cortical dynamics during and after a single session of left DLPFC piTBS in healthy young men. Frontal theta power between stimulation trains followed different trajectories during active and sham piTBS. The direct comparison of pre-to-post TEP changes identified no significant clusters, and the exploratory frontal and posterior N100 findings warrant independent confirmation. Active piTBS showed no clear benefit over sham on either behavioral measure. Tracking frontal theta power between trains across repeated piTBS sessions could help determine whether the first-session response is associated with later changes in cortical reactivity or depressive symptoms.

## 8. Author contributions

Conceptualization: SXW, WMWC, JJZ, TFY, GSK; Methodology: SXW, WMWC, RRZ, JJZ; Investigation: SXW, WMWC, VWML, TTL; Data curation: SXW, WMWC, RRZ, PPQ, MJ, AWX, JJZ; Software: SXW, WMWC, TTL, RRZ, JJZ; Analysis: SXW, WMWC, RRZ, JJZ; Visualization: SXW, WMWC, PPQ, MJ, AWX; Validation: WMWC, TTL, RRZ, PPQ; Writing-original draft: SXW, WMWC, GSK; Writing-review & editing: SXW, WMWC, VWML, TTL, RRZ, PPQ, MJ, AWX, JJZ, TFY, YN, GSK; Funding acquisition, Resources, Supervision, Project administration: GSK.

## 9. Data availability statement

The data that support the findings of this study are available from the corresponding author upon reasonable request.

## 10. Acknowledgments

We appreciate the facilities support provided by the University Research Facility in Behavioral and Systems Neuroscience at The Hong Kong Polytechnic University. We also thank all participants for their time and effort.

## 11. Declaration of generative AI use

During the preparation of this manuscript, the authors used Gemini 3 for language editing and proofreading purposes. The authors take full responsibility for the integrity and accuracy of all scientific content, including all data, analyses, interpretations, and conclusions presented in this work.

---

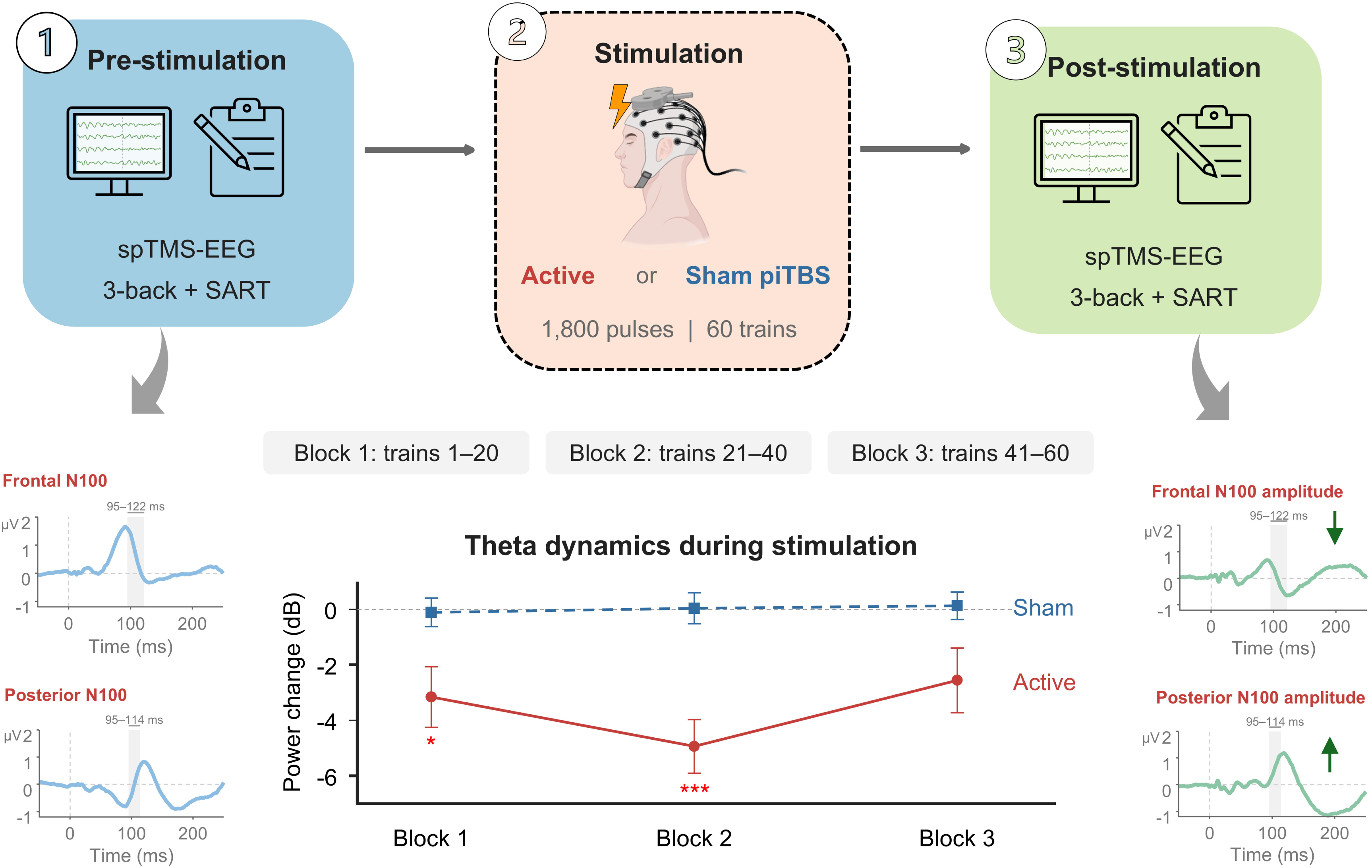

