## Supplementary Materials for "Acute Effects of Prolonged Intermittent Theta Burst Stimulation on Cortical Excitability: A Randomized Crossover TMS-EEG Study"

**Supplementary methods**

**S1. Electroencephalography**

EEG data were acquired using a 64-channel EEG cap with TMS-compatible Ag/AgCl electrodes (Easycap, Germany) arranged according to the international 10-20 system. The cap was connected to a DC-coupled amplifier (SynAmps2, USA). The electrodes FCz and AFz served as the online recording reference and ground, respectively. EEG signals were recorded and digitized at a sampling rate of 5 kHz with CURRY 7 software (Compumedics NeuroScan, USA). Electrode impedances were regularly checked and kept below 5 kΩ throughout the experimental sessions. All EEG recordings were conducted in a quiet, dimly lit room. Throughout EEG recording, participants remained in a seated position with their eyes closed and were instructed to stay awake. During the spTMS-EEG blocks, participants listened to white noise through intra-auricular earphones to limit the influence of auditory artifacts induced by the TMS clicks. The sound level was individually adjusted until the spTMS clicks at 100%RMT were masked by the white noise.

S1.1 **Preprocessing of piTBS-EEG data**

EEG data recorded during piTBS were analyzed using custom scripts in MATLAB R2024b (MathWorks, USA) and EEGLAB [1]. Continuous EEG data were first resampled to 1,000 Hz, high-pass filtered at 1 Hz, and notch filtered at 50 Hz. The data were then segmented into non-overlapping epochs from 1,000 to 7,000 ms relative to the final pulse of each of the 60 two-second piTBS trains, yielding 60 epochs per participant per condition. In addition, one baseline epoch was extracted for each session from the 6,000 ms period preceding the first pulse of the piTBS session. After epoching, noisy channels were identified and removed prior to independent component analysis. Independent component analysis was then performed, and artifactual components related to eye movements, muscle activity, cardiac activity, line noise, and channel noise were removed. Previously removed channels were then reconstructed using spherical interpolation, and the cleaned EEG data were band-pass filtered from 1 to 100 Hz and rereferenced to the common average. Power spectral density (PSD) was estimated using Welch’s method, and spectral power was extracted for the delta, theta, alpha, and beta frequency bands. To assess changes across the stimulation session, the 60 post-stimulation epochs were divided into three consecutive blocks of 20 epochs each (block 1: epochs 1-20; block 2: epochs 21-40; block 3: epochs 41-60). For each block, band power was averaged across epochs and normalized to baseline using a decibel transform [2], calculated as

$$\text{Power}_{\text{dB}}\text{=10}\text{log}_{\text{10}} \left( \frac{\text{Block Power}}{\text{Baseline Power}} \right)$$

Accordingly, positive and negative decibel values indicated relative increases and decreases in oscillatory power, corresponding to event-related synchronization (ERS) and desynchronization, respectively [3] (Figure 2B).

S1.2 **Preprocessing of spTMS-EEG data**

spTMS-EEG data were analyzed offline using version 2 of the fully automated advanced artifact removal for automated TMS-EEG processing (AARATEP) pipeline implemented in MATLAB R2024b (MathWorks, USA) (Figure 2A). spTMS-EEG data were epoched from -800 to +1000 ms relative to the TMS onset. The high-amplitude direct pulse artifact (-2 to +12 ms) was gated and reconstructed via autoregressive (AR) interpolation, fitting AR models to 20 ms segments flanking the artifact window to generate forward and backward predictions combined via sigmoidal-weighted blending to preserve per-pulse signal integrity. Signals were downsampled to 1000 Hz and baseline-corrected (-500 to -10 ms). To prevent the phase distortion and temporal smearing of low-latency stimulation transients into surrounding epochs, a modified AR-based piecewise high-pass filter (> 1 Hz) was implemented. This stage involved fitting AR models to data 100 ms prior and 30 ms after an extended artifact window (-6 to +36 ms) to extrapolate 500 ms segments forward and backward; the 1 Hz Butterworth high-pass filtering was applied separately to the pre- and post-pulse data, and the filter-induced signal alterations were sigmoidally blended across the artifact timespan and applied to the original non-extrapolated data. Artifact-laden channels were automatically flagged via a Wiener noise estimate (threshold = 10) and reconstructed using spherical spline interpolation. Early ocular artifacts were isolated and excised via an initial independent component analysis (ICA) coupled with automated ICLabel classification [4]. Source-space non-neuronal noise was suppressed using the spatial overdetermined noise unfolding algorithm (λ = 10^-1.5^) [5]. To mitigate sensor charge-decay artifacts, the dominant low-latency artifact topography was extracted from trial-averaged data (12–22 ms) and modeled using an exponential decay function (αe^−x/τ^), under mathematical constraints explicitly set to safeguard topographically overlapping, late-latency TEP components. Line noise was attenuated using a 48–52 Hz bandstop filter. A secondary ICA-ICLabel stage eliminated residual physiological noise based on empirical classification thresholds (eye score > 0.08, muscle score > 0.2, brain score < 0.3), a high-frequency amplitude ratio (> 8) within the 11–30 ms window to capture late muscle activity, and an absolute peak amplitude threshold (> 15 μV). Data were low-pass filtered (< 100 Hz) using a Butterworth filter and transformed to a common average reference. Pipeline source code is available at [github.com/chriscline/AARATEPPipeline](https://github.com/chriscline/AARATEPPipeline).

**S2. 3-back task**

The visual 3-back task was used to evaluate working memory, given its established sensitivity to prefrontal cortex function and its frequent use as a behavioral index of DLPFC-dependent cognitive processes [6]. Stimuli comprised uppercase letters (A–Z) presented pseudo-randomly in black (Consolas font, size 30) against a white background. Each letter was displayed for 500 ms, followed by a 3,000-ms inter-stimulus interval featuring a blank screen. Participants were required to press ‘1’ on the keyboard if the current letter matched the one presented exactly three trials prior (target condition), and ‘2’ if it did not match (non-target condition). In instances where participants pressed a button multiple times within a single trial, the reaction time (RT) was recorded based on the final keystroke. The task was administered in three consecutive blocks, with short rest periods provided between blocks. Each block contained 30 trials (10 targets and 20 non-targets). To properly establish the working memory buffer, the first three trials of each block served as fillers to initiate the 3-back sequence; these initial trials were strictly excluded from all subsequent behavioral data analyses, yielding 27 valid test trials per block. Task performance was assessed using *d′* (d-prime), a measure of sensitivity derived from the hit rate and false alarm rate: *d′* = Z (hit rate)−Z (false alarm rate).

**S3. Sustained Attention to Response Task**

The SART was administered to provide a complementary assessment of sustained attention and response inhibition as indices of prefrontal-dependent executive function [7]. The SART is approximately 4.3 min in duration and consists of 225 trials. Each trial is 1,150 ms in length. During the first 250 ms of each trial, either a Go stimulus, represented by the numbers ranging from 1 to 9 excluding 3, or a No-Go stimulus, represented by the number 3, is presented, followed by a 900-ms mask. Number stimuli are presented white on a black background with Arial Font randomly sized at either 4%, 7% 10%, 13%, or 16% of the displayable screen. Number stimuli are presented randomly. Participants respond using the space bar on their computer, and the response time for each trial is recorded. The ratio of Go to No-Go stimuli is 8 to 1 or 88.9% (200 of 225) Go to 11.1% (25 of 225) No-Go. Task performance was assessed using commission error rate. Commission error rate quantifies failures of response inhibition and is calculated as the proportion of No-Go stimuli on which participants made a response.

**S4. Statistical analysis**

S4.1 **piTBS-induced oscillatory responses**

PSD-derived alpha- and theta-band power values in a left frontal region of interest, defined a priori as the mean of electrodes F3 and AF3, were analysed because of the proximity of these electrodes to the stimulation target over the left dorsolateral prefrontal cortex. Alpha- and theta-band activity was selected based on prior literature indicating that frontal low-frequency oscillations are relevant to TMS-related cortical responses [8,9]. Separate rmANOVAs were performed for each frequency band, with condition (active piTBS vs. sham piTBS) and block (block 1, block 2, block 3) as within-subject factors. Sphericity was assessed using Mauchly’s test, and Greenhouse–Geisser correction was applied when sphericity was violated. Significant main effects or interactions were followed by Bonferroni-corrected post hoc paired-samples t-tests. One-sample t-tests against 0 dB were also conducted to assess deviations from baseline, with Bonferroni correction applied for multiple comparisons.

S4.2 **TMS-evoked potentials**

TEPs were analyzed using spatiotemporal cluster-based permutation tests at the global scalp level. Predefined time windows were used for each TEP component of interest: (N45 [30–50 ms], P60 [50–70 ms], N100 [95–125 ms] and P180 [160–200 ms]). These components are commonly described in the literature [10–13]. Within each component window, two-tailed dependent-samples *t*-tests were performed across all electrodes and time points in a Monte Carlo cluster-based permutation framework. Neighboring electrodes were defined using a distance-based criterion. Clusters were formed using a sample-level threshold of *p* < 0.05, quantified with the maximum cluster-sum statistic, and required a minimum of two neighboring electrodes. Cluster significance was determined from 5,000 random permutations. Within each component window, post-minus-pre TEP change waveforms were compared between active and sham piTBS. Exploratory analyses compared pre- and post-stimulation waveforms separately within each intervention condition. For each contrast, two-sided cluster *p* values were Bonferroni-adjusted across the four component windows. Adjusted *p* values below .05 were considered significant. For each significant cluster identified in the exploratory within-condition analyses, mean TEP amplitude was calculated by averaging across the electrodes and time points within that cluster. These values were entered into separate exploratory two-way repeated-measures ANOVAs (rmANOVAs) with the within-subject factors condition (active piTBS vs. sham piTBS) and time (pre-stimulation vs. post-stimulation). Main effects of condition and time, and the condition × time interaction were examined. Exploratory simple-effect comparisons assessed pre-to-post amplitude changes within each condition. The resulting p values were Bonferroni-adjusted.

S4.3 **Cognitive tasks**

Linear mixed-effects models were used to analyze d′ and commission error rate, with time point (pre-stimulation vs. post-stimulation), condition (sham piTBS vs. active piTBS), the condition × time interaction, and visit order (visit 1 vs. visit 2) included as fixed effects. Age and years of education were entered as continuous covariates. Participant was included as a random intercept to account for repeated measures. The significance of fixed effects was evaluated using Type III *F*-tests with Satterthwaite-approximated degrees of freedom. Post hoc comparisons of estimated marginal means were Bonferroni-adjusted.

**Supplementary figure**

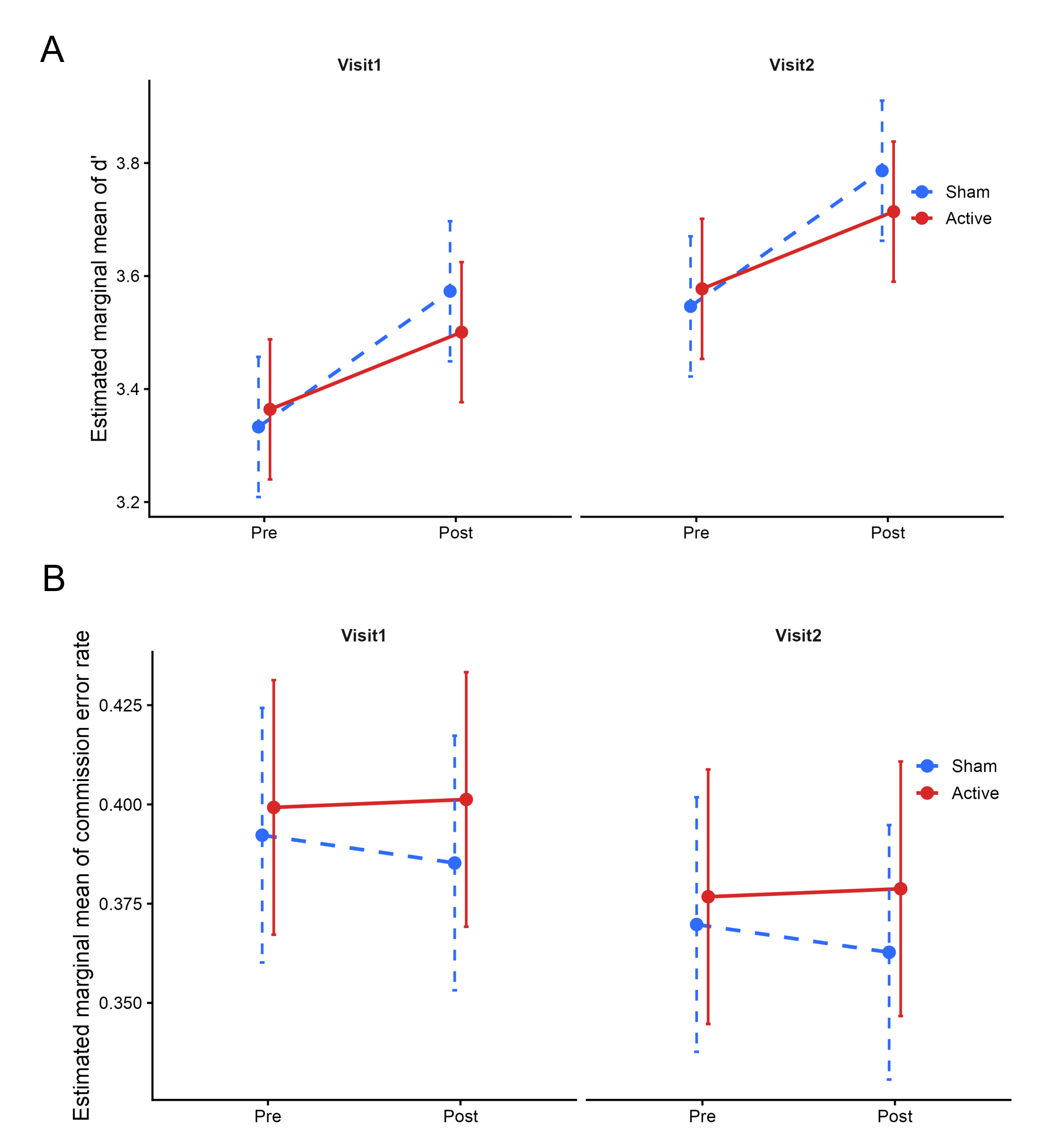

**Figure S1. Model-estimated task performance by time, piTBS condition and visit order.** (A) Estimated marginal means of 3-back discriminability (d′). (B) Estimated marginal means of SART commission error rate. Points represent estimated marginal means from the linear mixed-effects models, and error bars indicate standard errors. Lines connect pre- and post-stimulation estimates within each condition. Red solid lines indicate active piTBS, and blue dashed lines indicate sham piTBS. Visit 1 and Visit 2 are shown in separate subpanels. Estimates were adjusted for age and years of education.
